## Supplementary Tables 1 and 2 for "*In silico* logical modelling to uncover cooperative interactions in cancer"

Wild type, RPTP_L=1

| RPTP_L | HGF | EGF | ECM | TGFB | IL6 | DELTA | ROS | WNT | Existing phenotypes | Reachability probability | #Successful runs | #Runs |
|---|---|---|---|---|---|---|---|---|---|---|---|---|
| 1 | 0 | 0 | 0 | 0 | 0 | 0 | 0 | 0 | 20 | 1.0000 | N/A | N/A |
| 1 | 1 | 0 | 0 | 0 | 0 | 0 | 0 | 0 | 20 | 1.0000 | N/A | N/A |
| 1 | 0 | 1 | 0 | 0 | 0 | 0 | 0 | 0 | 20 | 1.0000 | N/A | N/A |
| 1 | 0 | 0 | 1 | 0 | 0 | 0 | 0 | 0 | 20 | 0.6262 | | |
| 1 | 0 | 0 | 1 | 0 | 0 | 0 | 0 | 0 | 23 | 0.1805 | 1.00E+05 | ### |
| 1 | 0 | 0 | 1 | 0 | 0 | 0 | 0 | 0 | 03 | 0.1933 | | |
| 1 | 0 | 0 | 0 | 1 | 0 | 0 | 0 | 0 | 01 | 1.0000 | N/A | N/A |
| 1 | 0 | 0 | 0 | 0 | 1 | 0 | 0 | 0 | 21 | 0.4695 | 1.00E+05 | ### |
| 1 | 0 | 0 | 0 | 0 | 1 | 0 | 0 | 0 | 01 | 0.5305 | | |
| 1 | 0 | 0 | 0 | 0 | 0 | 1 | 0 | 0 | 20 | 1.0000 | N/A | N/A |
| 1 | 0 | 0 | 0 | 0 | 0 | 0 | 1 | 0 | 02 | 1.0000 | N/A | N/A |
| 1 | 0 | 0 | 0 | 0 | 0 | 0 | 0 | 1 | 20 | 1.0000 | N/A | N/A |

NOTCH E1, RPTP_L=1

| RPTP_L | HGF | EGF | ECM | TGFB | IL6 | DELTA | ROS | WNT | Existing phenotypes | Reachability probability | #Successful runs | #Runs |
|---|---|---|---|---|---|---|---|---|---|---|---|---|
| 1 | 0 | 0 | 0 | 0 | 0 | 0 | 0 | 0 | 20 | 1.0000 | N/A | N/A |
| 1 | 1 | 0 | 0 | 0 | 0 | 0 | 0 | 0 | 20 | 1.0000 | N/A | N/A |
| 1 | 0 | 1 | 0 | 0 | 0 | 0 | 0 | 0 | 20 | 1.0000 | N/A | N/A |
| 1 | 0 | 0 | 1 | 0 | 0 | 0 | 0 | 0 | 20 | 0.6210 | 99999 | ### |
| 1 | 0 | 0 | 1 | 0 | 0 | 0 | 0 | 0 | 03 | 0.3790 | | |
| 1 | 0 | 0 | 0 | 1 | 0 | 0 | 0 | 0 | 01 | 1.0000 | N/A | N/A |
| 1 | 0 | 0 | 0 | 0 | 1 | 0 | 0 | 0 | 01 | 1.0000 | N/A | N/A |
| | | | | | | | | | | | N/A | N/A |
| 1 | 0 | 0 | 0 | 0 | 0 | 1 | 0 | 0 | 20 | 1.0000 | N/A | N/A |
| 1 | 0 | 0 | 0 | 0 | 0 | 0 | 1 | 0 | 02 | 1.0000 | N/A | N/A |
| 1 | 0 | 0 | 0 | 0 | 0 | 0 | 0 | 1 | 20 | 1.0000 | N/A | N/A |

Wild type, RPTP_L=0

| RPTP_L | HGF | EGF | ECM | TGFB | IL6 | DELTA | ROS | WNT | Existing phenotypes | Reachability probability | #Successful runs | #Runs |
|---|---|---|---|---|---|---|---|---|---|---|---|---|
| 0 | 0 | 0 | 0 | 0 | 0 | 0 | 0 | 0 | 20 | 1.0000 | N/A | N/A |
| 0 | 1 | 0 | 0 | 0 | 0 | 0 | 0 | 0 | 12 | 0.9370 | 1.00E+05 | ### |
| 0 | 1 | 0 | 0 | 0 | 0 | 0 | 0 | 0 | 02 | 0.0630 | | |
| 0 | 0 | 1 | 0 | 0 | 0 | 0 | 0 | 0 | 12 | 0.8191 | N/A | N/A |
| 0 | 0 | 1 | 0 | 0 | 0 | 0 | 0 | 0 | 02 | 0.1809 | N/A | N/A |
| 0 | 0 | 0 | 1 | 0 | 0 | 0 | 0 | 0 | 20 | 0.6081 | 99997 | ### |
| 0 | 0 | 0 | 1 | 0 | 0 | 0 | 0 | 0 | 03 | 0.3919 | | |
| 0 | 0 | 0 | 0 | 1 | 0 | 0 | 0 | 0 | 02 | 1.0000 | N/A | N/A |
| 0 | 0 | 0 | 0 | 0 | 1 | 0 | 0 | 0 | 21 | 0.4701 | 1.00E+05 | ### |
| 0 | 0 | 0 | 0 | 0 | 1 | 0 | 0 | 0 | 01 | 0.5299 | | |
| 0 | 0 | 0 | 0 | 0 | 0 | 1 | 0 | 0 | 20 | 1.0000 | N/A | N/A |
| 0 | 0 | 0 | 0 | 0 | 0 | 0 | 1 | 0 | 02 | 1.0000 | N/A | N/A |
| 0 | 0 | 0 | 0 | 0 | 0 | 0 | 0 | 1 | 20 | 1.0000 | N/A | N/A |

NOTCH E1, RPTP_L=0

| RPTP_L | HGF | EGF | ECM | TGFB | IL6 | DELTA | ROS | WNT | Existing phenotypes | Reachability probability | #Successful runs | #Runs |
|---|---|---|---|---|---|---|---|---|---|---|---|---|
| 0 | 0 | 0 | 0 | 0 | 0 | 0 | 0 | 0 | 20 | 1.0000 | N/A | N/A |
| 0 | 1 | 0 | 0 | 0 | 0 | 0 | 0 | 0 | 02 | 1.0000 | N/A | N/A |
| | | | | | | | | | | | N/A | N/A |
| 0 | 0 | 1 | 0 | 0 | 0 | 0 | 0 | 0 | 02 | 1.0000 | N/A | N/A |
| | | | | | | | | | | | N/A | N/A |
| 0 | 0 | 0 | 1 | 0 | 0 | 0 | 0 | 0 | 20 | 0.6017 | 99982 | ### |
| 0 | 0 | 0 | 1 | 0 | 0 | 0 | 0 | 0 | 03 | 0.3983 | | |
| 0 | 0 | 0 | 0 | 1 | 0 | 0 | 0 | 0 | 02 | 1.0000 | N/A | N/A |
| 0 | 0 | 0 | 0 | 0 | 1 | 0 | 0 | 0 | 01 | 1.0000 | N/A | N/A |
| | | | | | | | | | | | N/A | N/A |
| 0 | 0 | 0 | 0 | 0 | 0 | 1 | 0 | 0 | 20 | 1.0000 | N/A | N/A |
| 0 | 0 | 0 | 0 | 0 | 0 | 0 | 1 | 0 | 02 | 1.0000 | N/A | N/A |
| 0 | 0 | 0 | 0 | 0 | 0 | 0 | 0 | 1 | 20 | 1.0000 | N/A | N/A |

Wild type, HGF=1 and ECM=1

| RPTP_L | HGF | EGF | ECM | TGFB | IL6 | DELTA | ROS | WNT | Existing phenotypes | Reachability probability | #Successful runs | #Runs |
|---|---|---|---|---|---|---|---|---|---|---|---|---|
| 1 | 1 | 0 | 1 | 0 | 0 | 0 | 0 | 0 | 20 | 0.6226 | | |
| 1 | 1 | 0 | 1 | 0 | 0 | 0 | 0 | 0 | 23 | 0.1808 | 1.00E+05 | ### |
| 1 | 1 | 0 | 1 | 0 | 0 | 0 | 0 | 0 | 03 | 0.1966 | | |
| 0 | 1 | 0 | 1 | 0 | 0 | 0 | 0 | 0 | 03 | 1.0000 | N/A | N/A |

Wild type, EGF=1 and ECM=1

| RPTP_L | HGF | EGF | ECM | TGFB | IL6 | DELTA | ROS | WNT | Existing phenotypes | Reachability probability | #Successful runs | #Runs |
|---|---|---|---|---|---|---|---|---|---|---|---|---|
| 1 | 0 | 1 | 1 | 0 | 0 | 0 | 0 | 0 | 20 | 0.6249 | | |
| 1 | 0 | 1 | 1 | 0 | 0 | 0 | 0 | 0 | 23 | 0.1798 | 1.00E+05 | ### |
| 1 | 0 | 1 | 1 | 0 | 0 | 0 | 0 | 0 | 03 | 0.1953 | | |
| 0 | 0 | 1 | 1 | 0 | 0 | 0 | 0 | 0 | 03 | 1.0000 | N/A | N/A |

**Table S1: The EMT logical model predicts cooperation between RPTP_L, microenvironment signals and NOTCH E1.** Probabilities of reaching EMT phenotypes (columns existing phenotypes), according to the model read-outs AJ and FA, starting from an E1 phenotype, when RPTP_L is fixed at 1 (upper left table), or RPTP_L is fixed at 0 (middle left panel), or HGF and ECM are fixed at 1 (lower left table), or RPTP-L is fixed at 1 in the presence of a NOTCH E1 mutation (upper right table), or RPTP-L is fixed at 0 in the presence of a NOTCH E1 mutation (middle right table), or EGF and ECM are fixed at 1 (lower right table). Green colors denote when the levels of the microenvironmental inputs is set to 1.

| # | | UN | | M1 | | M2 | | M3 | | 10 | | 11 | | H2 | | 13 | | E1 | | H1 | | 22 | | H3 | |
|---|---|---|---|---|---|---|---|---|---|---|---|---|---|---|---|---|---|---|---|---|---|---|---|---|---|
| | | Sgle | Dble | Sgle | Dble | Sgle | Dble | Sgle | Dble | Sgle | Dble | Sgle | Dble | Sgle | Dble | Sgle | Dble | Sgle | Dble | Sgle | Dble | Sgle | Dble | Sgle | Dble |
| | NOTCH E1 | | NA | | NA | | NA | | NA | | NA | | NA | | NA | | NA | | NA | | NA | | | NA | |
| 1 | AKT KO | | | | | | | | | | | | | | | | | | | | | | | | |
| 2 | AKT E1 | | | | | | | | | | | | | | | | | | | | | | | | |
| 3 | BCatAJ KO | | | | | | | | | | | | | | | | | | | | | | | | |
| 4 | BCatAJ E1 | | | | | | | | | | | | | | | | | | | | | | | | |
| 5 | BCat KO | | | | | | | | | | | | | | | | | | | | | | | | |
| 6 | BCat E1 | | | | | | | | | | | | | | | | | | | | | | | | |
| 7 | CK1 KO | | | | | | | | | | | | | | | | | | | | | | | | |
| 8 | CK1 E1 | | | | | | | | | | | | | | | | | | | | | | | | |
| 9 | CSL KO | | | | | | | | | | | | | | | | | | | | | | | | |
| 10 | CSL E1 | | | | | | | | | | | | | | | | | | | | | | | | |
| 11 | DVL KO | | | | | | | | | | | | | | | | | | | | | | | | |
| 12 | DVL E1 | | | | | | | | | | | | | | | | | | | | | | | | |
| 13 | ECadAJ E2 | | | | | | | | | | | | | | | | | | | | | | | | |
| 14 | ECadAJ KO | | | | | | | | | | | | | | | | | | | | | | | | |
| 15 | ECadAJ E1 | | | | | | | | | | | | | | | | | | | | | | | | |
| 16 | ECad KO | | | | | | | | | | | | | | | | | | | | | | | | |
| 17 | ECad E1 | | | | | | | | | | | | | | | | | | | | | | | | |
| 18 | EGFR KO | | | | | | | | | | | | | | | | | | | | | | | | |
| 19 | EGFR E1 | | | | | | | | | | | | | | | | | | | | | | | | |
| 20 | ERK KO | | | | | | | | | | | | | | | | | | | | | | | | |
| 21 | ERK E1 | | | | | | | | | | | | | | | | | | | | | | | | |
| 22 | FAK_SRC KO | | | | | | | | | | | | | | | | | | | | | | | | |
| 23 | FAK_SRC E1 | | | | | | | | | | | | | | | | | | | | | | | | |
| 24 | FAK_SRC E12 | | | | | | | | | | | | | | | | | | | | | | | | |
| 25 | FAT4 KO | | | | | | | | | | | | | | | | | | | | | | | | |
| 26 | FAT4 E1 | | | | | | | | | | | | | | | | | | | | | | | | |
| 27 | GSK3B KO | | | | | | | | | | | | | | | | | | | | | | | | |
| 28 | GSK3B E1 | | | | | | | | | | | | | | | | | | | | | | | | |
| 29 | HGFR KO | | | | | | | | | | | | | | | | | | | | | | | | |
| 30 | HGFR E1 | | | | | | | | | | | | | | | | | | | | | | | | |
| 31 | HIF1a KO | | | | | | | | | | | | | | | | | | | | | | | | |
| 32 | HIF1a E1 | | | | | | | | | | | | | | | | | | | | | | | | |
| 33 | ILK KO | | | | | | | | | | | | | | | | | | | | | | | | |
| 34 | ILK E1 | | | | | | | | | | | | | | | | | | | | | | | | |
| 35 | ITG_AB KO | | | | | | | | | | | | | | | | | | | | | | | | |
| 36 | ITG_AB E1 | | | | | | | | | | | | | | | | | | | | | | | | |
| 37 | JAK KO | | | | | | | | | | | | | | | | | | | | | | | | |
| 38 | JAK E1 | | | | | | | | | | | | | | | | | | | | | | | | |
| 39 | JNK KO | | | | | | | | | | | | | | | | | | | | | | | | |
| 40 | JNK E1 | | | | | | | | | | | | | | | | | | | | | | | | |
| 41 | LATS KO | | | | | | | | | | | | | | | | | | | | | | | | |
| 42 | LATS E1 | | | | | | | | | | | | | | | | | | | | | | | | |
| 43 | MEK KO | | | | | | | | | | | | | | | | | | | | | | | | |
| 44 | MEK E1 | | | | | | | | | | | | | | | | | | | | | | | | |
| 45 | miR200 KO | | | | | | | | | | | | | | | | | | | | | | | | |
| 46 | miR200 E1 | | | | | | | | | | | | | | | | | | | | | | | | |
| 47 | NFkB KO | | | | | | | | | | | | | | | | | | | | | | | | |
| 48 | NFkB E1 | | | | | | | | | | | | | | | | | | | | | | | | |
| 49 | p120AJ KO | | | | | | | | | | | | | | | | | | | | | | | | |
| 50 | p120AJ E1 | | | | | | | | | | | | | | | | | | | | | | | | |
| 51 | PAK KO | | | | | | | | | | | | | | | | | | | | | | | | |
| 52 | PAK E1 | | | | | | | | | | | | | | | | | | | | | | | | |
| 53 | PI3K KO | | | | | | | | | | | | | | | | | | | | | | | | |
| 54 | PI3K E1 | | | | | | | | | | | | | | | | | | | | | | | | |
| 55 | RAF1 KO | | | | | | | | | | | | | | | | | | | | | | | | |
| 56 | RAF1 E1 | | | | | | | | | | | | | | | | | | | | | | | | |
| 57 | RAP1 KO | | | | | | | | | | | | | | | | | | | | | | | | |
| 58 | RAP1 E1 | | | | | | | | | | | | | | | | | | | | | | | | |
| 59 | RAS KO | | | | | | | | | | | | | | | | | | | | | | | | |
| 60 | RAS E1 | | | | | | | | | | | | | | | | | | | | | | | | |
| 61 | RPTP KO | | | | | | | | | | | | | | | | | | | | | | | | |
| 62 | RPTP E1 | | | | | | | | | | | | | | | | | | | | | | | | |
| 63 | SLUG KO | | | | | | | | | | | | | | | | | | | | | | | | |
| 64 | SLUG E1 | | | | | | | | | | | | | | | | | | | | | | | | |
| 65 | SMAD KO | | | | | | | | | | | | | | | | | | | | | | | | |
| 66 | SMAD E1 | | | | | | | | | | | | | | | | | | | | | | | | |
| 67 | SNAIL KO | | | | | | | | | | | | | | | | | | | | | | | | |
| 68 | SNAIL E1 | | | | | | | | | | | | | | | | | | | | | | | | |
| 69 | STAT3 KO | | | | | | | | | | | | | | | | | | | | | | | | |
| 70 | STAT3 E1 | | | | | | | | | | | | | | | | | | | | | | | | |
| 71 | TCF_LEF KO | | | | | | | | | | | | | | | | | | | | | | | | |
| 72 | TCF_LEF E1 | | | | | | | | | | | | | | | | | | | | | | | | |
| 73 | TGFBR KO | | | | | | | | | | | | | | | | | | | | | | | | |
| 74 | TGFBR E1 | | | | | | | | | | | | | | | | | | | | | | | | |
| 75 | YAP-TAZ KO | | | | | | | | | | | | | | | | | | | | | | | | |
| 76 | YAP-TAZ E1 | | | | | | | | | | | | | | | | | | | | | | | | |
| 77 | ZEB KO | | | | | | | | | | | | | | | | | | | | | | | | |
| 78 | ZEB E1 | | | | | | | | | | | | | | | | | | | | | | | | |

**Table S2: Phenotypes compatible with the 80 LoF and GoF single mutants and by the 78 combinations of double mutants involving a NOTCH GoF mutation.** Each column corresponds to phenotypes reached by each single LoF (KO) and GoF (E1) mutant indicated on the left, combined (Dble) or not (Sgle) with the NOTCH E1 mutation Grey boxes indicate phenotypes retrieved by the model under the specified condition and input configuration. For some perturbations, the phenotypic landscapes are reshaped, leading to the appearance of the novel unnamed phenotypes 10, 11, 13 and 22.
